# Selective Recruitment of Cortical Neurons by Electrical Stimulation

**DOI:** 10.1101/213017

**Authors:** Maxim Komarov, Paola Malerba, Paul Nunez, Eric Halgren, Maxim Bazhenov

## Abstract

Despite its critical importance in experimental and clinical neuroscience, at present there is no systematic method to predict which neural elements will be activated by a given stimulation regime. Here we develop a novel approach to model the effect of cortical stimulation on spiking probability of neurons in a volume of tissue, by applying an analytical estimate of stimulation-induced activation of different cell types across cortical layers. We utilize the morphology and properties of axonal arborization profiles obtained from publicly available anatomical reconstructions of the twelve main categories of neocortical neurons to derive the dependence of activation probability on cell type, layer and distance from the source. We then propagate this activity through the local network incorporating connectivity, synaptic and cellular properties. Our work predicts that (a) intracranial cortical stimulation induces selective activation across cell types and layers; (b) superficial anodal stimulation is more effective than cathodal at cell activation; (c) cortical surface stimulation focally activates layer I axons, and (d) an optimal stimulation intensity exists capable of eliciting cell activation lasting beyond the end of stimulation.

## Introduction

Brain stimulation is widely used to probe the properties of neural systems (Halgren et al. 1978; Tehovnik and Lee 1993; Britten and van Wezel 1998; Graziano et al. 2002), to normalize dysfunction (e.g., deep brain stimulation for Parkinsonian symptoms (Blumenfeld and Brontë-Stewart 2015; Baizabal-Carvallo and Alonso-Juarez 2016; Papageorgiou et al. 2016) and epileptic patients (Nagaraj et al. 2015), Direct-Current Stimulation for stroke patients (Hummel and Cohen 2006)), or to manipulate brain activity, including enhancing memories and learning (Fröhlich et al. 2015; Prehn and Flöel 2015; Savic and Meier 2016; Summers et al. 2016). While the practice is widespread and scalable, its application is limited by the difficulty of predicting which cells (if any) are going to spike due to an input, and which specific synaptic mechanisms are going to be recruited and modulated by a given stimulation protocol (Draganski et al. 2014). In addition, the secondary effects of the directly activated neurons on other cells may be more distributed and prolonged than the direct effects themselves. Furthermore, depending on the goal of stimulation, the effect of interest could be driving cells to spike or inducing sub-threshold changes. Stimulation can synchronize (Polanía et al. 2011), de-synchronize (Popovych and Tass 2012), excite (McIntyre et al. 2004; Rahman et al. 2013) and/or suppress (McIntyre *et al.* 2004; Rahman *et al.* 2013) neuronal activity. However, it has been difficult to infer these observations by applying the physics of current fields to cortical anatomy and physiology.

Considerable work has attempted to define the specifics of current flow when the cortex is directly stimulated (Meffin et al. 2012; Draganski *et al.* 2014). In particular, anatomical measures derived from brain scans, cell reconstructions, and other measures have been used to populate finite element models, leading to patient-specific suggestions regarding where current would flow for a given electrode placement(Datta et al. 2009; Datta et al. 2011). Such models are often “passive”, meaning the active properties of neurons (e.g., spike generation, synaptic dynamics, intrinsic currents) are not taken into account.

The effects of stimulation have been examined using electrophysiological and/or optical methods. However, the results have been ambiguous due to limitations in recording from an entire cortical volume with high temporal and spatial resolution (Histed et al. 2009; Tehovnik and Slocum 2013). This underscores the need for careful modeling to predict plausible outcomes that can be verified separately in different layers and cell types. Furthermore, an empirically validated modeling approach could be extended easily to a variety of contemplated electrode configurations and stimulation regimes.

In this work, we develop a method that estimates the effect of extracellular electrical stimulation on the different classes of neocortical neurons. Our approach begins with calculation of the activation function (Rattay 1987; Rattay and Aberham 1993; Rattay 1999), which represents the stimulation-induced effective current across neuronal membranes. Our approach is novel, in that it convolves the activation function with axonal and dendritic arborizations of different cell types, obtained from public databases of over five hundred morphological reconstructions of twelve different classes of cortical neurons, while also accounting for myelination properties, morphological variability within each cell class, and the different orientations and positions of neurons. We apply this method to predict the direct activation probabilities of cortical cell types across all layers for short (~200 *μs*) current pulse typical of clinical stimulation protocols. We then calculate the consequences of this direct activation by propagating activation through a multi-layer neocortical network model with realistic channel and synaptic elements, to predict the net effect of stimulation on network activity. Our work predicts (a) that intracranial cortical stimulation induces selective activation across cell types and layers; (b) cathodal vs anodal stimulation have distinct effects on different cell types; (c) cortical surface stimulation focally activates layer I axons, and (d) the existence of an optimal stimulation intensity capable of eliciting cell activation lasting beyond the end of stimulation.

## Materials and Methods

### Estimating electric potential: a single electrode

Below we assume the current delivered by the stimulation electrode *(I)* a fixed amount. We consider a Cartesian coordinate system *(X,Y,Z)* centered at the electrode, where the *Z* axis is oriented perpendicular to the electrode surface and goes in depth across cortical layers; the axes X and Y are oriented parallel to the surface and electrode plate (see Fig.1a). To first approximation, we assume that the current *I* across the insulating layer of the electrode is uniform over the electrode surface. Each infinitesimally small element of the electrode *dS* then produces a current source

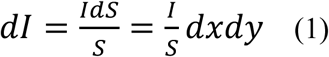

Where *S* is the electrode surface, *dS = dxdy* is the infinitesimal electrode element. Here, the pair *(x, y)* locates the current source on the surface of the circular electrode (where *Z*=0). The elemental current source produces a tissue potential at distance *L* from the source given by

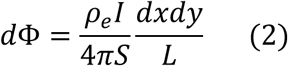

where *ρ*_*e*_ is the resistivity of the tissue, *S* = *A*^2^ is the surface area of the electrode, *A* is the size of the electrode and *I* is the total current generated by the electrode.

Note that the distance can be expressed as 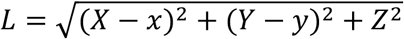.

Hence, the potential in the Cartesian coordinate system *(X,Y,Z)* is given by

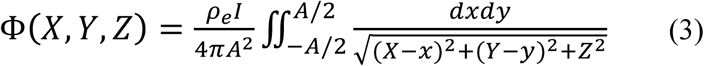

The integral form of the equation (3) is due to the need to compute the contribution of the entire electrode surface area *A* as a source of electrical current. However, at large distances the finite size of the electrode has a nominal effect on the electrical field. Indeed, the rapid decay of potential with respect to distance from the source is evident. Note that for tissue locations further away from the electrode plate, the point-source approximation provides the following simplified expression to compute the electrical potential field, (see Methods, Estimating electric potential: a single electrode):

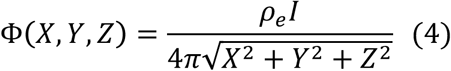

This estimate well approximates our exact solution (in equation (1)) at a distance from the electrode, however it fails for locations right below the electrode plate, where the finite size of the electrode should be taken into account (Fig. 1c). This distinction is particularly relevant when considering the potential field in cortical layers I and II, immediately below the surface, which are also the most affected by stimulation due to the decay of Φ with depth. Hence, we emphasize that close to the electrode, the point-source approximation does not yield an estimate that can be used to predict precisely the effect of cortical stimulation.

### Estimating activation function

According to one-dimensional cable theory the dynamics of transmembrane voltage of axonal segments can be computed as follows:

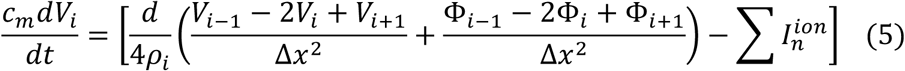

here *V*_*i*_1_, *V*_*i*_, *V*_*i*+1_ denote transmembrane voltages of the neighboring axonal compartments (subindex denotes number of the compartment), *c*_*m*_ = 1*μF*/*cm*^2^ is a capacitance of membrane per square unit area, d stands for diameter of the axon (typically between 10 *μm* and 1 *μm*), *ρ*_*i*_ = 300Ω ⋅ *cm* is a resistivity of axoplasm, Δ*x* is a discretization parameter, which defines length of the compartment. The term 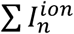 describes the sum of intrinsic ionic current such as fast potassium and sodium currents for spike generation, leak currents and others. As one can see, the effective transmembrane current, which arises due to extracellular electrical stimulation is described by the term

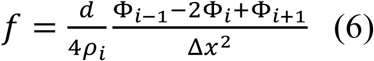

where Φ_*i*_,Φ_*i-i*_,Φ_*i+i*_ stand for extracellular potentials in the vicinity of axonal compartments. In fact, *f* represents activation function (Rattay 1987) and in the limit Δ*x*→ 0 can written as 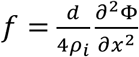 (here axis x represents direction of axonal fiber). We use activation function (computed along reconstructed axons) to estimate probability of axonal activation. In order to calculate activation function, we estimated the direction of the axonal segments using the position of neighboring compartments up to 10 microns apart from the point where direction was calculated. In doing so, we effectively reduce the impact that jitter in the edges of the reconstruction can have over the numerical calculation of the activation function.

### Selecting cells reconstructions within available databases

In order to obtain cell reconstructions, we use publicly available resources for neuronal morphologies (Ascoli et al. 2007). Table 1 summarizes the datasets used in our analysis.

**Table 1.**
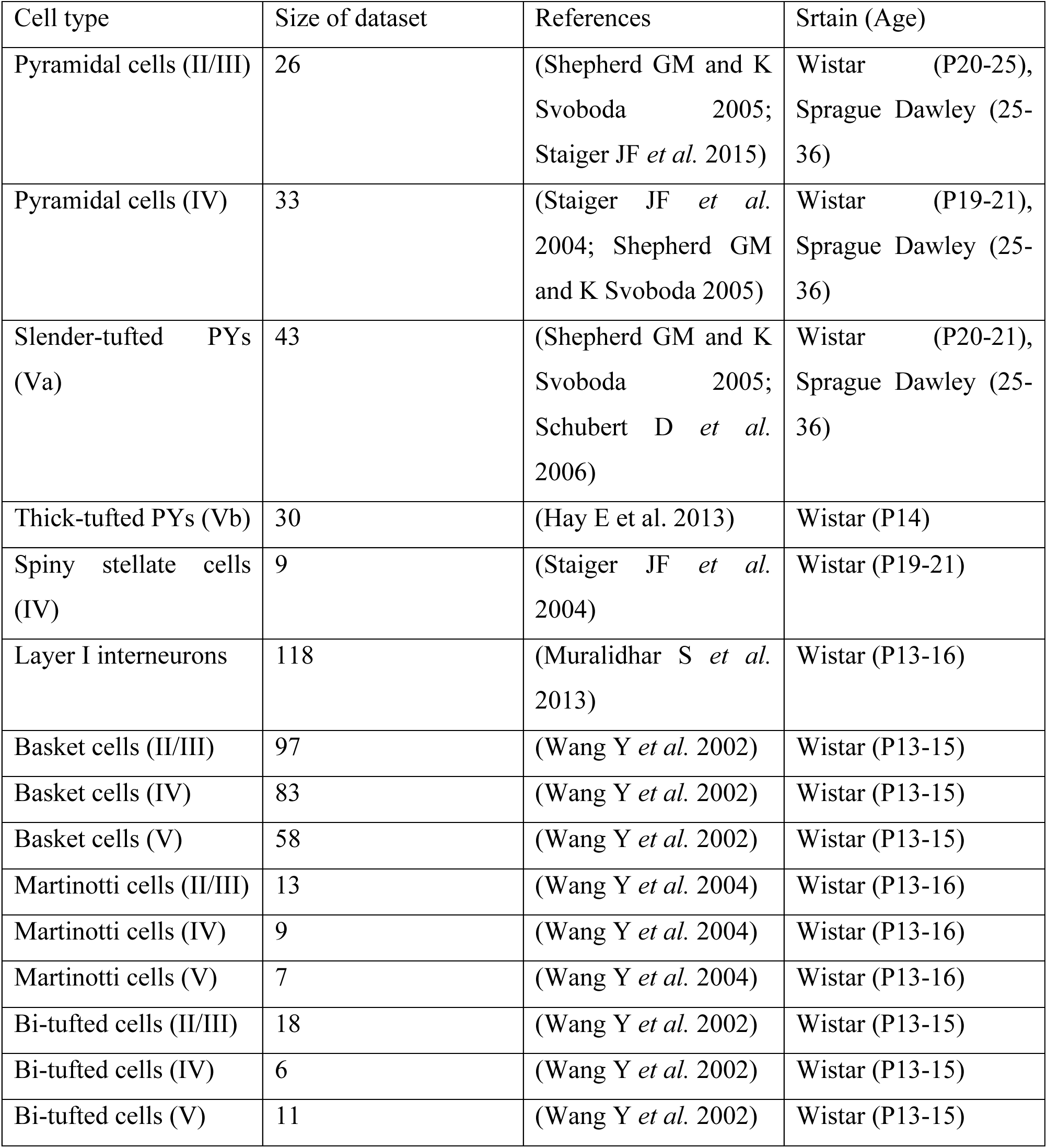
Summary of datasets with reconstructed cells.

All cell reconstructions were corrected for tissue shrinkage and aligned when necessary. The overall dataset is not homogenous, since cells were obtained from distinct experiments, which used rats of different age (ranges from P13-P36). However, for every cell type the parameters of the probability calculations were adjusted in order to account for possible differences in thickness of the cortex and cell size. Using data provided in the references (see Table 1), we estimated approximate boundaries for each layer, and used those values in our statistical analysis to predict activation probability. The variations introduced turned out to be small or negligible (see also comparative analysis in (Ramaswamy and Markram 2015)), and would not affect our main results.

### Computing the average axonal arborization for a given cell type

Averaged axonal density (Fig.2B) represents overall morphological properties of a given type of neurons (among those in Table 1) and gives the general intuition on how a given cell type can be affected by electrical stimulation. The axonal density was computed as follows: first, all 3D reconstructions of cells of a certain type were aligned in space and centered at soma, such that axis coordinates *(x, y, z)* correspond to width, height and depth of the slice respectively. Second, all reconstructions were superimposed in a 3d volume, and the axonal density was constructed on a grid of a size 100 points in the x direction, by 100 points in the *y* direction, by 50 points in the z direction. Finally, the 3D density array was averaged across the z-axis and the result is plotted in Fig.2B in a logarithmic scale.

### Computing the averaged probability of activation

Our computations of the averaged probability of activation are based on the calculation of the activation function and the trigger area. The trigger area comprises axonal segments with sufficiently high (above *3 pA/μm*^2^) values of activation function, which can initiate axonal action potential in unmyelinated segments of axons (e.g. nodes of Ranvier). The threshold was defined based on comparison to experimentally recorded current-distance relation for pyramidal cells (see section Results for details). Note that this threshold was computed for axon initial segment, but we used the same value for nodes of Ranvier. For unmyelinated fibers we assumed 5-fold larger threshold, since their excitability is lower due smaller concentration of sodium channels. To compute the averaged probability for a certain class of cells (Table 1) located at distance R_0_ from the electrode we applied the following steps:

a. We took one of the anatomical reconstructions and placed it at a distance R_0_ from the electrode at a certain depth Z within the layer that the cell belongs to.
b. We computed activation function *f* for axonal segments of the reconstructed cell. Next, the function was evaluated against the threshold (*3 pA/μm^2^* for myelinated and 60 *pA/μm^2^* for unmyelinated fibers). The segments, which possess large (above threshold) activation function were marked as a trigger area, whose elements may initiate axonal response.
c. We then transformed length of the trigger area *L* into probability of spiking. For myelinated fibers, the trigger area should contain at least one node of Ranvier to initiate axonal response. To find an activation probability, we discretize the trigger area into small segments of length *k=1 μm* which is a typical length of a node of Ranvier. Note that the probability that a given segment is a node of Ranvier can be approximated as a ratio *p*_*n*_ = *k/D* where D denotes mean internodal distance (we used 100 *μm* in all estimations). Next, the probability that axonal fiber of length *L*does not contain any nodes nodes of Ranvier can be approximated as (1 – *p*_*n*_)^*N*^. Here *N*= *Lk* denotes the number of of segments of length *k* that can fit into a fiber of total length *L*. Hence, the overall probability p of response for myelinated axon can be estimated as 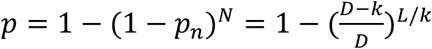 For unmyelinated fibers, whose entire membrane is exposed to extracellular space, we assumed binary dependence of probability on *L*: any *L*>0 (presence of trigger area) produced activation, while absence of trigger area (*L*=0) meant no activation.
d. The steps a-c were repeated for various locations (different Z) and for various orientations of the cell. Note that cells tend to grow towards the surface and occupy a significant area in layer I (Ramaswamy and Markram 2015). Therefore, in our calculations the depth coordinate Z varied in the range [*Z*_min_, *Z*_min_ + *cL*_*s*_], where *Z*_*min*_ is a minimal possible depth of the soma, *L*_*s*_ is a layer size and *c* is a coefficient (0.1 for all cells except basket cell, for which *c*=0.4). The minimal depth for a cell soma *Z*_*min*_ is given by the upper boundary of the cell’s layer for cell types that are compact (e.g. basket cells in infragranular layers, whose arborization does not reach the surface). For large cells, which show arborizations that can reach the cortical surface (e.g. all pyramidal cells), the *Z*_*min*_ value is taken as the length from the soma to the highest point in ascending arbor, whether such branch is axonal or dendritic.
e. Combining the results from the above step we have an overall probability that a given cell reconstruction would be activated.
f. Steps (a) to (d) were repeated for every cell reconstruction from the pool of available neurons. The probability was calculated averaging across all cells, to represent the overall likelihood of axonal activation for a given cell class at distance R_0_ from the electrode.

### Computational model of the canonical cortical circuit: rationale, equations and parameters

#### Rationale

The network model represents a cortical column, which contains pyramidal (PY), spiny stellate (SC), basket (BC) and Martinotti (MC) cells from layers II-V (Fig. 7A). Note that there is a large diversity of different types of interneurons in the cortex (Markram et al. 2004), but we restrict ourselves to the most common (Markram *et al.* 2004) and the most important types in the context of our study. BCs constitute about 50% of all interneurons in the cortex and form a major source of lateral inhibition within the layers (Markram *et al.* 2004). MCs are likely to be activated in all layers due to their specific form of axonal arborization (Fig.2, Fig. 4). Moreover, MCs comprise a significant part of all interneurons in infragranular layers (Markram *et al.* 2004; Wang et al. 2004) where they specifically target excitatory cells in layer IV (Wang *et al.* 2004).

Excitatory neurons (PYs and SCs) (Schubert et al. 2003) and inhibitory MCs (Markram *et al.* 2004) were modeled as regular spiking cells with spike rate adaptation (Fig. 5B, top voltage trace shown for PYs), whereas inhibitory BCs were modeled as fast spiking cells (Fig. 5B, middle voltage trace). All interneurons had lower leak current, which resulted in a higher responsiveness in comparison to the excitatory cells (Povysheva et al. 2006). Cell dynamics was governed by Hodgkin-Huxley-type kinetics, which includes fast Na^+^-K^+^ spike generating mechanism (for all types of cells), high-threshold activated Ca^2+^ current (for PY and SCs) and slow calcium-dependent potassium (AHP) current (for regular spiking cells).

#### Equations

The membrane potential is governed by the following equation (Traub et al. 1991):

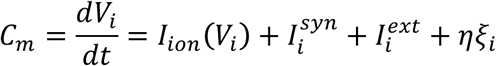

The ionic currents *I*_*ion*_(*V*_*i*_), which are responsible for intrinsic cells dynamics, read:

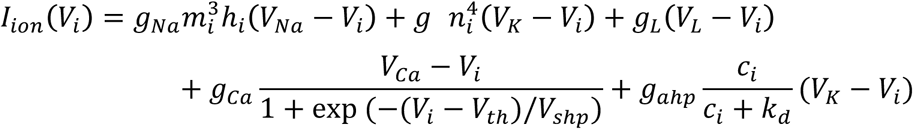

The gating variables *m*_*i*_, *n*_*i*_, *h*_*i*_ evolve according to:

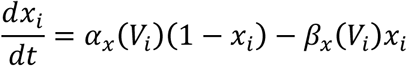

where *x*_*i*_ is one of gating variables. The functions *α*_*x*_(*V*) and *β*_*x*_(*V*) are:

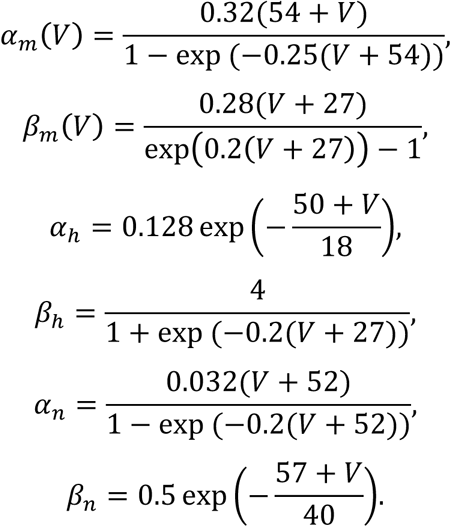

Calcium concentration *c*_*i*_ obeys the following equation:

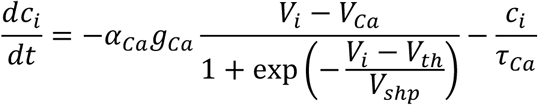

and governs calcium-dependent hyperpolarizing potassium current 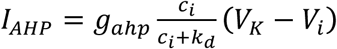, which is responsible for spike-frequency adaptation. The synaptic input was modeled according to:

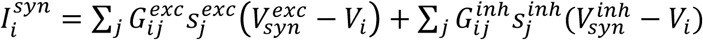

where synaptic variables 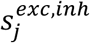 are governed by the following equation:

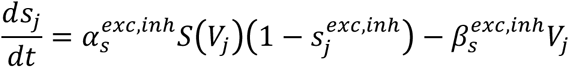

The function *S*(*V*) reads:

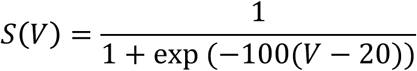

The term 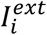 denotes applied external current (stimulus that leads cells to spike during stimulation), and the term *ηξ*_*i*_(*t*) corresponds to the fluctuations in the afferent input (representing spontaneous background activity) which are given by a white noise process with the following properties: <*ξ*_*i*_(*t*) >= 0, <*ξ*_*i*_(*t*)*ξ*_*i*_(*t*-*t*_0_) >=*δ*(*t*-*t*_0_). Here *δ(t)* stands for Dirac delta function, constant *η* determines the strength (standard deviation) of the noise. All the model parameters are listed below (unless specified in the description of simulations):

Cells were synaptically coupled by AMPA-type excitatory and GABA_a_-type inhibitory connections. The structure of synaptic connections represented a random graph. The strength and probability of connections depended on the layer and the cell type, resembling structure of a canonical cortical circuit (Thomson et al. 2002). Each layer contained relatively strong recurrent excitatory (PY->PY and PY->BC) and lateral inhibitory (BC->PY) connections. According to canonical architecture, PYs within layers II/III were driven by strong connections from layers IV and Va(Thomson *et al.* 2002; Shepherd and Svoboda 2005; Staiger et al. 2015). Excitatory cells from layers IV and Va (slender-tufted PY) were strongly coupled through recurrent excitatory connections within the layers(Schubert *et al.* 2003; Staiger et al. 2004; Schubert et al. 2006). In addition, PYs within layer Va received moderate excitatory input from layer IV(Schubert *et al.* 2006).

Inhibitory lateral connections from interneurons to PY were more intense and strong than excitatory ones(Holmgren et al. 2003). BCs formed local connections that project to excitatory cells within their own layers(Markram *et al.* 2004). In accordance with experimental studies(Wang *et al.* 2004), MCs from layer IV projected to excitatory cells from layer IV and II/III, whereas layer V MCs contacted excitatory cells from layer IV.

#### Model Parameters

The following summarizes all parameters of the individual cells and network architecture used in the model (Traub *et al.* 1991):

**A.** Parameters that were common across all cell types:

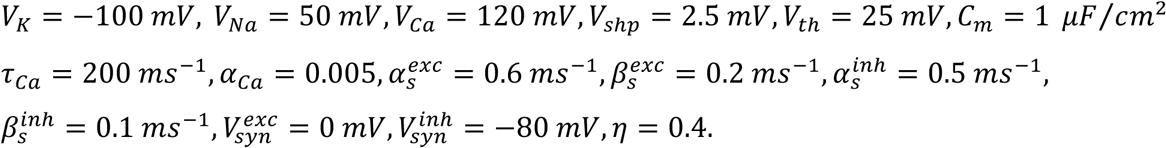

**B.** Specific parameters:

B.1 PYs and SCs:

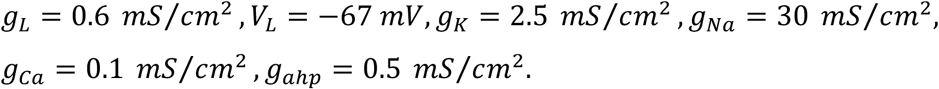

B.2 MCs:

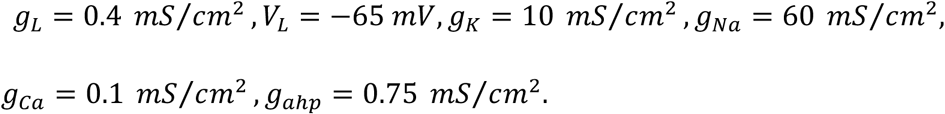

B.3 MCs:

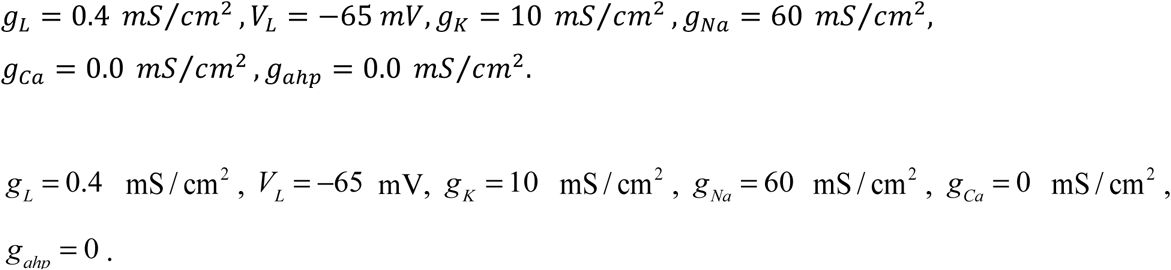

**C.** Network structure and connectivity:

Structure:

**Table 2.** Structure of the network. PY-pyramidal neurons, BC-basket cells, SC-excitatory spiny stellate cells, MC-Martinotti cells.

| Type of cortical cells | Number of cells |
| --- | --- |
| PY II/III | 200 |
| BC II/III | 50 |
| PY+SC IV | 200 |
| BC IV | 25 |
| MC IV | 25 |
| PY Va (slender) | 50 |
| BC V | 25 |
| MC V | 25 |

Connectivity:

**Table 3.** Connectivity within the network. PY-pyramidal neurons, BC-basket cells, SC-excitatory spiny stellate cells, MC-Martinotti cells

| To | From | Type | Strength | Probability |
| --- | --- | --- | --- | --- |
| PY, BC II/III | PY II/III | AMPA | 0.75 | 0.1 |
| PY II/III | PY, SC IV | AMPA | 1.5 | 0.05 |
| PY II/III | PY Va (slender) | AMPA | 0.75 | 0.05 |
| PY, SC, BC, MC IV | PY, SC IV | AMPA | 1.5 | 0.1 |
| PY, SC IV | PY II/III | AMPA | 0.25 | 0.05 |
| PY Va (slender) | PY Va (slender) | AMPA | 0.75 | 0.1 |
| BC, MC | PY Va (slender) | AMPA | 0.75 | 0.1 |
| PY II/III | BC II/III | GABA | 3 | 0.25 |
| PY II/III | MC IV,V | GABA | 0.5 | 0.05 |
| PY, SC IV | BC IV | GABA | 3 | 0.25 |
| PY, SC IV | MC IV | GABA | 3 | 0.25 |
| PY, SC IV | MC V | GABA | 2 | 0.2 |
| PY Va (slender) | BC V | GABA | 3 | 0.25 |
| PY Va (slender) | MC IV,V | GABA | 0.5 | 0.05 |

## Results

Our Results section is organized as following. We first provide estimate of the electric potential created by small surface located electrode in the tissue volume and we calculate activation function for neuronal fibers within cortical tissue. We next discuss what impact such activation has on specific populations of cortical neurons and we introduced a collection of the reconstructed cortical cells, together with the average profile of their axonal arborizations. We follow by computing a threshold to predict based on the activation function when applied stimulation is capable to trigger a response in the neuron of a given type and location. This allows us to calculate activation probability function, depending on the planar distance between a cell soma and the electrode, which is specific for different cell types. We then discuss the role of myelination in activation probability and we also ask how the probabilities of cell activation depend on the amplitude and polarity (anodal vs cathodal) of the stimulating current, and whether this dependence is similar across different cell types with different anatomies. Finally, we apply our analysis to predict network population response in canonical cortical circuit receiving surface cortical stimulation.

### Analytical estimate of the electric potential in tissue volume

**Fig.1.**
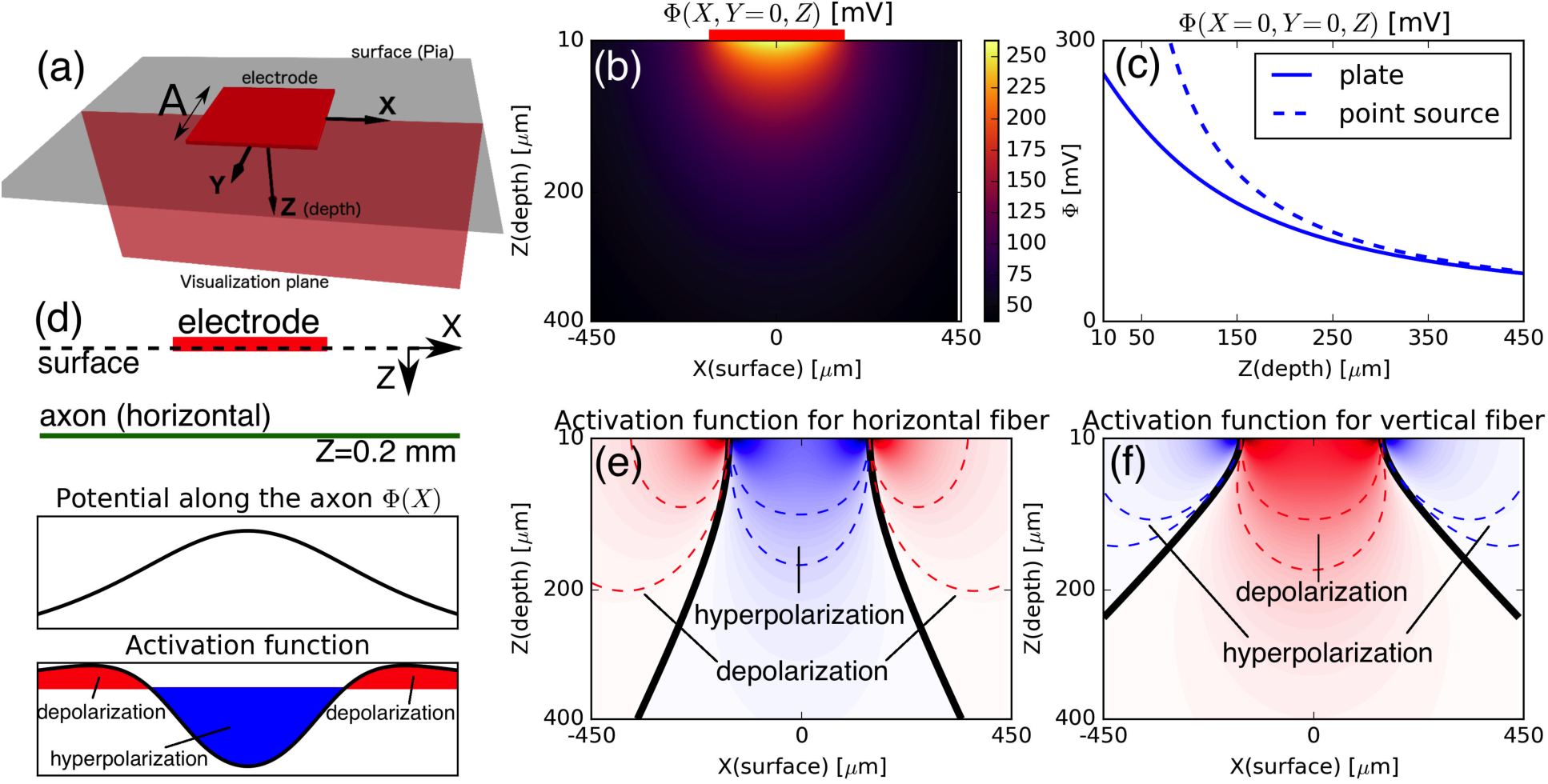
Electric potential under electrode plate and its effect on axonal fibers (the case of anodal stimulation is shown) (**a**) Schematic representation of the electrode in the coordinate system (X,Y,Z). Electrode is located on the surface (gray), center of the coordinate system corresponds to the center of the electrode. (**b**) Electric potential Φ(*X, Y,Z)* on the plane Y=0 (marked by red in panel (a)). (c) Comparison of the electric potential induced by point source (eq. (4), dashed curve) and finite-size square plate (eq. (3), solid curve) at varying depth Z and fixed X=Y=0. (d) Top panel shows schematic representation of the electrode (red) and horizontally oriented axon (green) on (X,Z) plane (Y=0). Bottom panels show potential Φ(x) and activation function 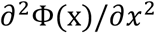 along axonal fiber (anodal stimulation). (e,f) Activation function for horizontally (e) and vertically (1) oriented fibers as a function of coordinates X,Z on the plane Y=0. Black solid curves separate areas of depolarization (red) and hyperpolarization (blue). Note that for cathodal stimulation the activation function is exactly opposite (area of depolarization and hyperpolarization are interchanged).

Below we consider a homogenous tissue under a single electrode (with the reference at infinity), which is used as a current source (our results also extend to stimulation applied by an array of electrodes). We considered the case of one electrode placed directly on the cortical surface (Fig. 1a), and electrode size and range of currents comparable to common experimental settings (Ha et al. 2016). For stimulation protocols that place the current source further from cortical surface, the current would have to pass through other tissues, such as skull, dura, arachnoid or pia maters before reaching the cortical layers. In such cases, our analysis would have to take into account potential capacitive properties, which effectively act as frequency filters, and current diffusion (Nunez and Srinivasan 2005). To estimate the effect of stimulation on the tissue, we first found the electrical field potential of our current source, based on the shape of the electrode and the total current injected into the tissue. Assuming that the current is uniform across the electrode surface, our source represents a homogeneous square electrode, and the resulting electric field potential can be calculated using the following expression (see Methods)

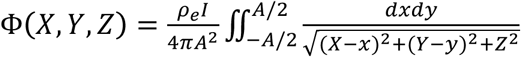

where *I* denotes net current, ρ_*e*_ is extracellular resistivity and A is the length of the square electrode edge (Fig.1a). In our analysis we use A=150 μm and net current I is in the range [0, 150] μA. For completeness, note that the integral in expression (1) has an analytical form (see Methods, Estimating electric potential: a single electrode). Fig. 1b shows the electric potential Φ, computed using equation (1), on the plane Y=0 (represented as a red plane in Fig. 1a).

The effect of applying a constant electric field to neuronal fibers can be described within one-dimensional cable theory, completed with the inclusion of the activation function (Rattay 1987, 1999). According to this theory, the activation function *f(X,Y,Z)* defines the effective transmembrane current that neuronal fibers (e.g. axons) receive due to extracellular electric stimulation. Technically, *f* can be computed as the second order spatial derivative of the electric potential Φ along neuronal fibers. For the simplest case of a perfectly straight axon, oriented horizontally in our Y=0 plane (Fig.1d), the activation function has the following form:

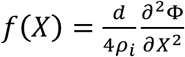

where *d* is the diameter of the axon, and ρ_*i*_ is the resistivity of the axoplasm. Hence, the shape of the electric potential Φ(*X*) along the axon (Fig.1d, middle panel) determines the function *f(X)* (Fig.1d, lower panel). As shown by Rattay (Rattay 1999), the activation function is a powerful tool for analyzing the effect of electrical stimulation on neuronal fibers, since it provides putative activation and suppression zones along the fibers. In case of a horizontal axon receiving anodal stimulation, the activation function along the axon shows that stimulation has a hyperpolarizing effect in the area right below the electrode (Fig.1d, blue area) and slightly depolarizing on portions of the fiber further to the sides (red areas). The distance of the axonal fiber from the electrode also influences the effect of stimulation, as shown in Fig 1e. The interplay of fiber orientation and placement in space is also shown in Fig. 1f, which presents the activation function *f(X,Y,Z)* for vertically (f) oriented axons. In Fig 1e-f, the color code emphasizes that the area below the electrode has a hyperpolarizing effect on horizontal fibers but a depolarizing effect on vertical ones. The spatial/orientation selectivity of the hyperpolarization/depolarization effect is still present when considering cathodal rather than anodal stimulation, with the caveat that, since the activation function would be reversed, the areas of depolarization/hyperpolarization would switch roles compared to Fig.1e-f.

### Neuronal response to stimulation can be predicted using reconstructed anatomy to define average arborization profiles

So far, we obtained an analytical estimate of the electric potential, and the activation function for neuronal fibers within cortical tissue. We next discuss what impact such activation has on specific populations of cortical neurons. The answer to this question depends on several factors including: (a) Effect of a current injected at a given distance from a neuron on the membrane voltage of that neuron; (b) Distribution of different neuron types across layers. Indeed, different neuron types are distributed differently across cortical layers(DeFelipe 2011), and it is legitimate to expect that their different properties and placement in the cortex would affect if and how these neurons respond to the surface electrical stimulation.

**Fig.2.**
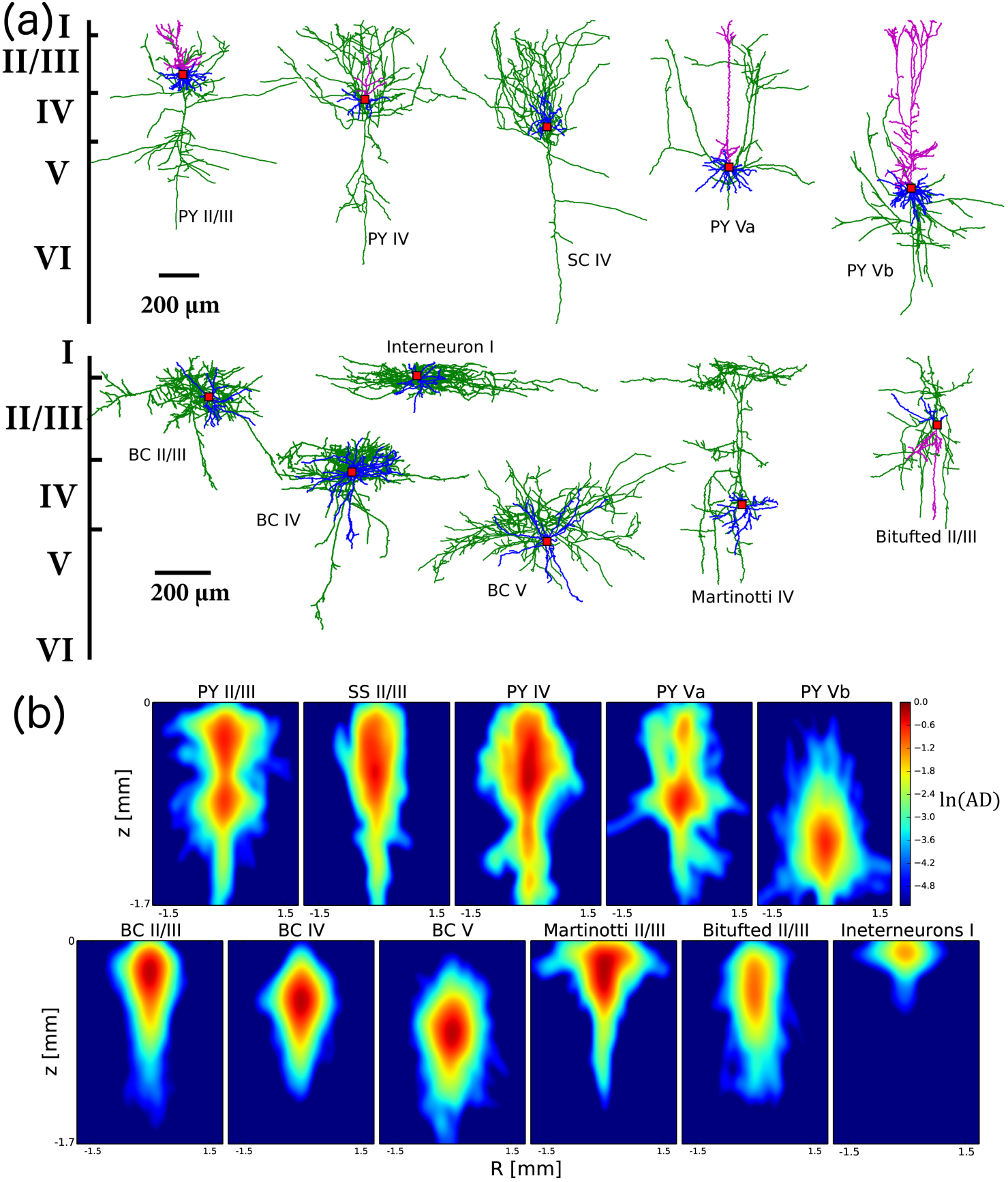
Anatomical reconstructions of the main types of cortical neurons. **A:** Typical anatomical profiles for the main types of cortical neurons. Green denotes axon, purple-apical dendrite, blue-basal dendrite, red dot shows soma position. Top row exhibits excitatory cells (PY-pyramidal neurons, SC-spiny stellate cell). Bottom row contains inhibitory interneurons (BC-basket cell). **B:** Averaged axonal densities formed by the neurons of each specific type. Color denotes logarithm of averaged axonal density (AD), computed over a set of available reconstructions of cortical cells. Logarithmic scale was used for better visualization of axonal arborization. This provides a general intuition on the generic shape of the axonal arborization for distinct types of cortical cells, which is crucial for the analysis.

A growing body of evidence supports the idea that electrical stimulation directly drives a response in a cell by triggering an action potential in nodes of Ranvier, or by activating the axon initial segment (orthodromic spikes) (Porter 1963; Stoney et al. 1968; Ranck 1975; Gustafsson and Jankowska 1976; Swadlow 1992; Nowak and Bullier 1998, 1998; Rattay 1999; Tehovnik et al. 2006). Thus, we concentrate our analysis on estimating the effects of stimulation on cells’ axonal fibers, and ignore their dendritic arborizations. To answer this question we need to study how, on average, the axonal arbors of different cell types are laid out in the cortical tissue. Reconstructing with any precision a specific and complete 3D volume of cortical tissue is yet impossible, and would introduce a strong limitation in our estimates by being too sensitive to the specifics of the very tissue reconstructed. Still, multiple databases containing the detailed reconstructions of different cell types from different preparations are available(Ascoli *et al.* 2007). Thus, we propose a new method to build for each cell type an approximation of a volume distribution of its axonal arborization, by taking advantage of the large datasets available on the specific anatomy of different cortical cell types.

Fig 2A shows an example of the reconstructions for the different cell types we considered in this study: pyramidal cells (PYs), excitatory spiny stellate cells (SCs) from layer IV, basket cells (BCs), Martinotti cells (MCs) and bi-tufted interneurons. There are two types of PYs in layer V: slender-tufted neurons from layer Va(Shepherd and Svoboda 2005; Schubert *et al.* 2006) and thick-tufted cells from layer Vb (Ramaswamy and Markram 2015). BCs include three subtypes according to the classification proposed in recent experimental works(Wang et al. 2002; Markram *et al.* 2004): large, nest and small basket cells. According to the canonical cortical microcircuit model(Douglas et al. 1989; Thomson *et al.* 2002; da Costa and Martin 2010; Defelipe et al. 2012), PYs and SCs from layers IV and Va receive input from thalamus and then innervate superficial layers, providing an incoming flow of information into cortical column (Shepherd and Svoboda 2005; Schubert *et al.* 2006). In turn, thick-tufted PYs (Vb) integrate the overall activity within and across columns, both neighboring and distant (Schubert et al. 2001; Schubert *et al.* 2006), and project their output to subcortical regions (Ramaswamy and Markram 2015).

Interneurons in cortex are very diverse in their morphology and functionality (Markram *et al.* 2004). BCs constitute about 50% of all inhibitory cells in cortex and form the primary source of lateral inhibition within layers, targeting somas and/or proximal dendrites of PYs (Wang *et al.* 2002). MCs comprise another significant fraction of interneurons, which can form cross-layer as well as cross-columnar inhibitory connections. These cells have a specific structure, with dense axonal arborization in layer I where they inhibit tuft and proximal dendrites of PYs from all layers (Wang *et al.* 2004). MCs are numerous especially in infragranular layers (Markram *et al.* 2004), where they are also known to specifically target basal dendrites of excitatory neurons from layer IV (Wang *et al.* 2004). In cortex, there are several other classes of interneurons which are found in fewer numbers: layer I interneurons, bipolar, double bouquet and bi-tufted cells (Markram *et al.* 2004; Muralidhar et al. 2013). Our analysis includes bi-tufted and layer I interneurons, as representative examples (in the context of our study) of this dendrite-targeting class of interneurons. Our population of reconstructed layer I interneurons contains 3 distinct subclasses (classification from (Muralidhar *et al.* 2013)): small, horizontal and descending interneurons.

Looking across multiple single-cell reconstructions for the same cell type, we can now design an average profile of the probability that a cell axonal arborization would occupy a given volume across layers (details on the procedure are introduced Methods). These averages are shown in Fig 2B, in logarithmic scale to emphasize the details of the differences across shapes. All types of excitatory cells, except thick-tufted PYs from layer Vb, have relatively dense axonal arborization in the top layer, which is reached by the strongest current density during surface-placed electrode stimulation (Fig.1). As for the interneurons, BCs axonal arborizations are largely contained within the layer occupied by their soma, while Martinotti and bi-tufted cells show axons with a wider vertical span and a large footprint in the top layer (Fig.2B). The axonal density of layer I cells is mainly confined in the top layer with small traces towards layer II, due to so-called descending interneurons (Muralidhar *et al.* 2013).

### Average axonal arborization and activation function provide an estimate for probability of axonal spiking response

In the previous sections we first found the electric potential in the tissue under the stimulating electrode and computed the activation function for axons (Fig 1). We then introduced a collection of the reconstructed cortical cells, together with the average profile of their axonal arborizations (Fig 2). To integrate the anatomical data and the estimated activation functions, we need to identify when the activation function is capable to trigger a response in the neuron. To compute such threshold, we matched the experimentally measured current-distance relationship leading to direct activation of cortical cells by the depth electrode (Stoney *et al.* 1968; Tehovnik *et al.* 2006).

**Fig.3.**
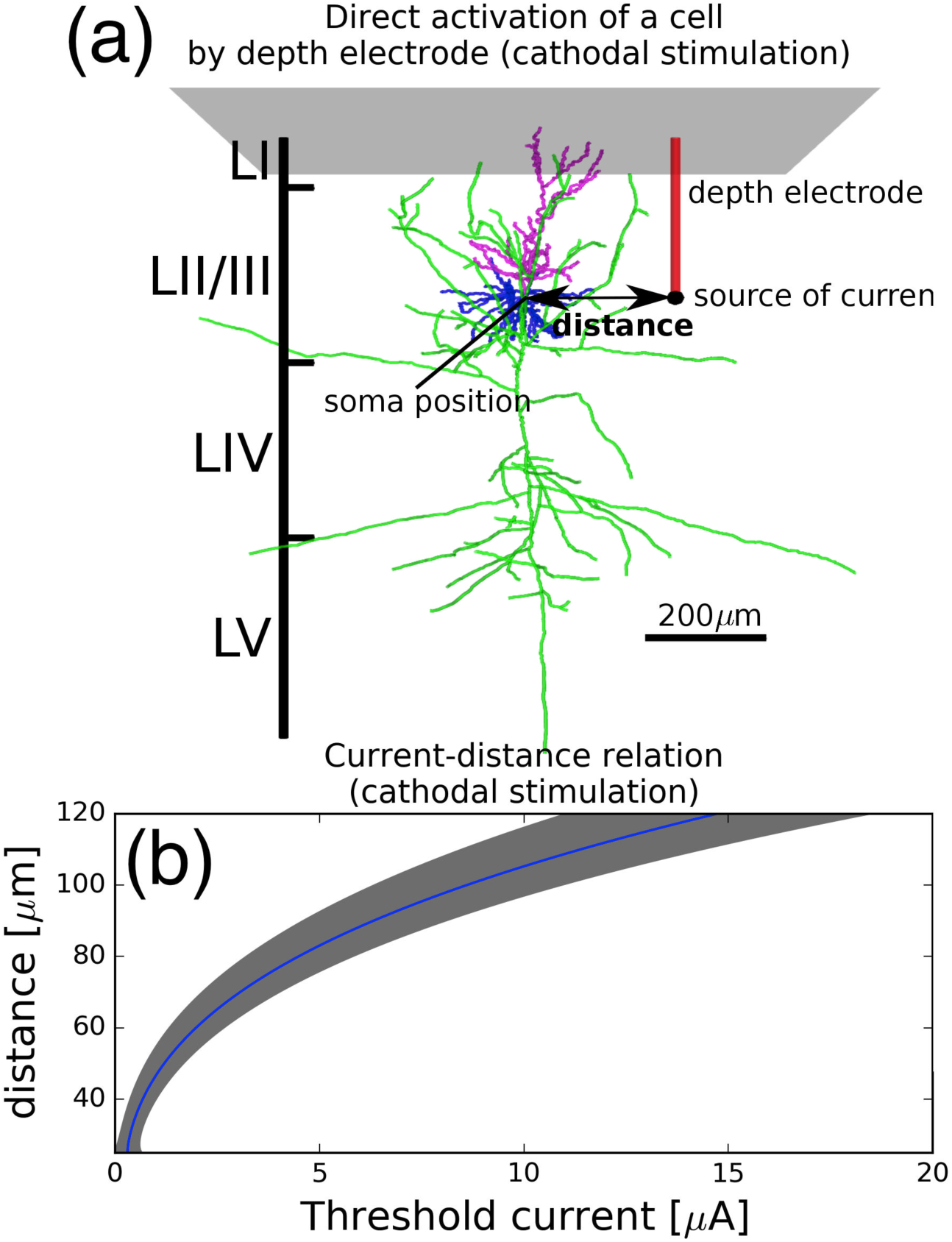
Theoretical method faithfully reproduces current-distance relation observed in experiments (Stoney SJ *et al*. 1968) (direct activation of cell by depth electrode). **(a)** Schematic representation of depth electrode (point source of current) and pyramidal cell. Green color denotes axons, purple and blue colors show apical and basal dendrites correspondingly. **(b)** Current-distance relation for direct activation of pyramidal cells (initial segment) by depth electrode. Blue curve represents average dependence across array of different reconstructions and rotations of pyramidal cells, gray area denotes mean plus/minus standard deviation. An ensemble of 15 cells was used.

Specifically, the experimental data we aim to match define a value of the threshold injected current *I*, which one has to apply to induce a threshold effective current *f* at the initial segment (located at distance *d* from the electrode). We used a 200μs duration stimulus pulse, typical of empirical *in vivo* microstimulation experiments (Stoney *et al.* 1968; Tehovnik *et al.* 2006). In Fig 3a, we show a representation of the *in vivo* experiment, which we mirrored in our model. Depth electrode (modeled as a point source of current) was placed near a cell body of a reconstructed pyramidal neuron from layer II/III. Using equations (2,3), we computed the activation current *f* at the axon initial segment (since the experimental data was focused on orthodromic activation), for different values of stimulation current *I* and distances *d*. We found that for all fixed values *f* = *Const* the resulting relation *I*(d) had a characteristic quadratic form, which qualitatively resembled experimental dependences of the threshold activation current on distance (Stoney *et al.* 1968; Tehovnik *et al.* 2006). By choosing *f* = *f*_*th*_ = 3 *pA*/*μm*^2^, we perfectly recovered the experimentally observed current-distance relation (Fig.3b, compare with Fig.8B in (Stoney *et al.* 1968)). Consistent with experimental data (Stoney *et al.* 1968; Ranck 1975; Tehovnik *et al.* 2006), we found that when the electrode was placed at a depth close to the soma (vertical coordinate in Fig.3a) we were able to find *f*_*th*_ only for cathodal current, while anodal current could not induce any depolarizing effect on the axonal initial segment. In what we discuss below, we will use this threshold value *f*_*th*_ to define the activation probability of axonal segments.

Once we estimated a threshold for each axonal segment, we can consider the overall effect of stimulation on a cell by taking into account its entire axonal arbor. To estimate the probability of cell activation we first computed the activation function *f* along the entire axon arbor and, by comparing *f* to the threshold *f*_th_, we identified which axonal segments were potentially activated (Fig.4, red markers). All together they form a total “trigger” area of length *L*. Thus, for the trigger area to initiate axonal activation in case of myelinated axons, it should contain at least one node of Ranvier where the current exceeds the threshold value *f*_th_. Therefore, we assume that the overall probability of cell activation depends on the length of the trigger area L and the probability of occurrence of nodes of Ranvier, which can be found using the average internodal distance (Ford et al. 2015) (see Materials & Methods for details). Intuitively, larger length of trigger area L and/or smaller internodal distance along the axon lead to a higher activation probability. In case of unmyelinated fibers, the entire membrane of axon is exposed to the extracellular space and, therefore, the cell activation probability depends only on the length L. For cell types with unmyelinated axons, we assumed a binary dependence: any L>0 (presence of trigger area) produced activation, while absence of trigger area (L=0) meant no activation. However, since unmyelinated axons are less excitable, their threshold of activation *(f*_th_) is much higher compared to nodes of Ranvier and axonal hillock: in our computations we used a threshold 20-fold larger. In summary, the length L of the trigger area can be computed using the activation function, and then transformed into probability of spiking for a given cell.

**Fig.4.**
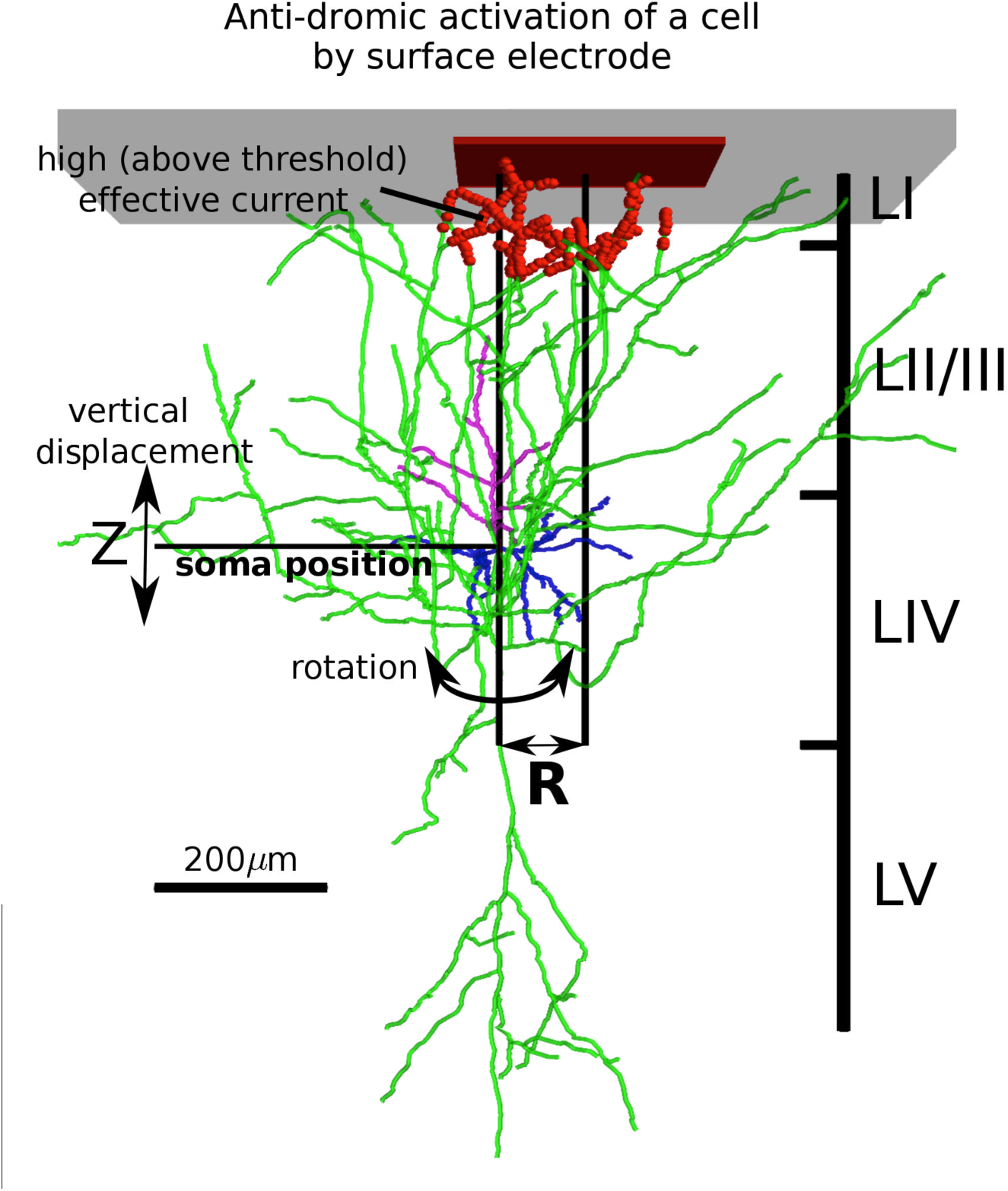
Estimation of the activation probability induced by surface stimulation. An example of typical layer IV pyramidal cell is shown. For each cell, we assigned **R**, and **Z** (depth) parameters. Activation function identifies its trigger area (red markers), where the effective current is above threshold. Action potentials can be initiated in these segments and propagate along the axonal arborization. To populate a statistical set (to find the average probability of spiking), each cell reconstruction was shuffled by rotating and shifting along the vertical axis (indicated by bold arrows), and multiple reconstructions were considered for each cell type (up to a total of 561 cells, see Table 1 in Methods: Selecting cell reconstructions within available databases).

In order to account for natural variability in cortical cell types and their locations with respect to the current source, we used the following approach, represented in Fig 4. For each anatomical reconstruction of a given cell type (up to a total of 561 cells, see Table 1 in Methods: Selecting cell reconstructions within available databases), we assigned a position R, which marked its planar distance from the center of the electrode plate, and a coordinate Z (depth), which located its soma within the appropriate cortical layer. We then identified (based on the length of trigger area L) probability of axonal activation for a given cell configuration within the cortical volume. Furthermore, we rotated the cell and shifted its soma in the vertical direction (for a range of Z values that still kept the cell within its type-defining layer, see Fig.4). As a result, we obtained numerous samples for a given neuron reconstruction placed at a distance R, and for each of them we evaluated if the neuron would be activated. The probability of activation for a given cell reconstruction (across all available rotations and vertical shifts) was given by the fraction of samples that were activated over the total number of samples. We repeated this procedure for each reconstructed cell belonging to a given cell type (see Table 1 in Materials and Methods), obtaining a probability of activation for each of them. We then considered the average of all these probabilities a faithful estimate of the probability of activation for a cell of a given type placed at distance R from the electrode.

### Distinct cell types show different profiles of activation probability

Our method defines an activation probability function, depending on the planar distance between a cell soma and the electrode (R in Fig 4), which can be different for different cell types. In Fig. 5 we present the results of our analysis applied to stimulation by a superficial cortical electrode. Since our simplified initial analysis suggests that anodal stimulation is most effective at depolarizing vertically oriented axonal arbors (see Fig 1E-F), consistently with general interpretation of experimental data (Ranck 1975), we present our analysis in the case of anodal stimulation. This section is organized into several parts, which discuss different cell types and the role of myelination in activation probability.

**Fig.5.**
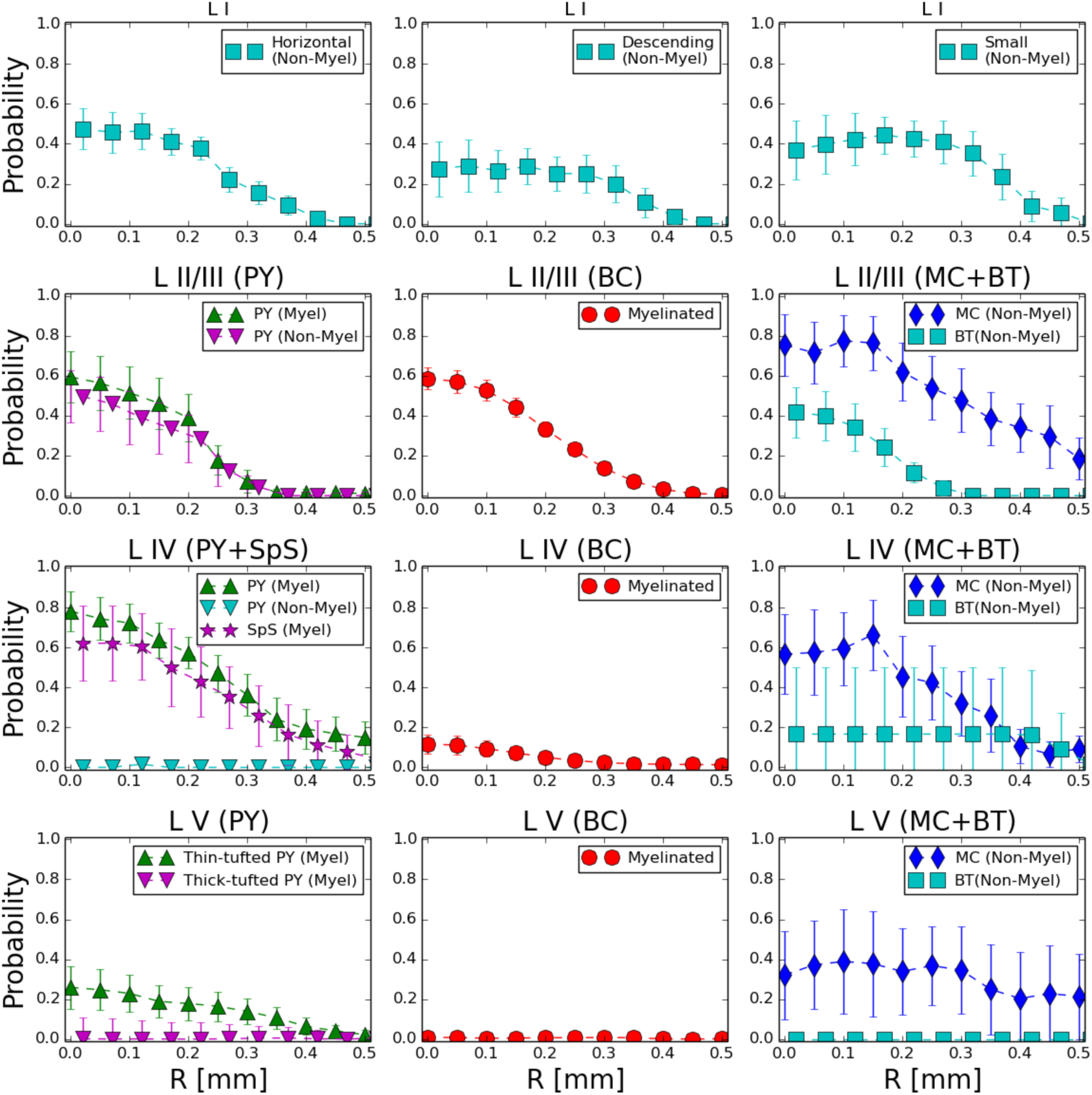
Probability of stimulation-induced activation is different across layers and cell types. The top row shows direct activation probability for 3 distinct types of layer I interneurons. Rows 2-4 (top to bottom) correspond to layers II-V. The left column contains probability for excitatory cells (pyramidal and spiny stellate), middle column contains data on soma/proximal dendrite-targeting interneurons (basket cells) and the right column contains probability for tuft/proximal dendrite-targeting interneurons (Martinotti and bitufted cells). The data represent anodal stimulation (*I* = 275 *μA*).

#### Excitatory cells

Our method predicts that pyramidal cells in most cortical layers would be only moderately activated by the superficial stimulation (Fig. 5, left column, rows 2-4), with the exception of excitatory cells (PYs and SCs) in layer IV, which have a fairly high probability to spike (80% right below the electrode) in response to the current stimulation. Since layer IV excitatory cells receive input from thalamus and other subcortical structures, and locally amplify such input (by strong recurrent connectivity) before projecting it to layer II/III pyramidal cells, their higher probability of activation in response to input suggests a possibility for superficial stimulation to be able to compensate for lacking subcortical inputs (for example following injury).

Within layer V, slender-tufted PYs (Va) have a moderate chance of direct activation (Fig.5, bottom row, left), compared to thick-tufted PYs, which do not seem to be directly recruited by the surface stimulation (Fig. 5, bottom row, left). This difference is consistent with their average axonal arborizations (Fig 2B): slender-tufted PYs tend to project their axons to the superficial layers (Shepherd and Svoboda 2005; Schubert *et al.* 2006), while thick-tufted PYs axonal density is sitting away from the superficial layers. Since thick-tufted PYs are the main output of a cortical column (Ramaswamy and Markram 2015), their activation effectively controls whether external electrical stimulation can influence downstream signaling to other brain regions. Hence, to activate the cortical output, external stimulation will need first to trigger enough of a local circuit response, so that the thick-tufted pyramidal cells in layer V can be recruited by the evoked neuronal activity of other cell types.

#### Basket cells

The central column of Fig. 5 (rows 2-4) shows that activation of BCs is very layer-specific; in particular, layer II/III BCs are easily activated, while deeper layer BCs are much less likely to be recruited. This estimate accounts for myelination in their axonal fibers (discussed in detail below), and is consistent with the localized organization of BCs average axonal densities (Fig 2B). Since BCs are the primary source of inhibition within each layer (they are the largest fraction of interneuron found in any cortical column (Markram *et al.* 2004)), their activation profile has the potential to shape the spiking within the cortical network. The fact that layer IV BCs are not directly recruited by input current further enables layer IV excitatory cells to trigger activity in the cortical column.

#### Other interneurons (Martinotti, Bitufted and layer I cells)

The rightmost column and top row of Fig. 5 describe the likelihood of activation for other non-parvalbumin interneuron types (Markram *et al.* 2004), which have unmyelinated axons and are likely to contact pyramidal cells in their distal dendrites. It is important to note that MCs from the infragranular layers (IV-VI) also target basal dendrites of the excitatory cells in layer IV(Wang *et al.* 2004). Our analysis shows that MCs have high activation probability in layers II/III and V, which is consistent with their specific axonal density distribution (Fig.2B). Extensive axonal arborization in layer I led to a high activation probability even though their axons are unmyelinated. Note that in our dataset, layer IV MCs show less activation than MCs in other layers, driven by the atypical shape of the axonal arbors in the reconstructions available.

Bitufted cells have moderate probability of activation only in supragranular layers. Expectedly, layer I interneurons being the closest to the current source, also exhibit high activation probability (Fig. 5, top row). This predicts that a surface stimulation would recruit a fair amount of spiking in cells that are responsible for diffused and cross-layer inhibitory signaling, within and across cortical columns. It may further suggest that cortical stimulation by the surface electrode, while capable of triggering spikes in excitatory cells in deep layers, is not likely to evoke very strong excitatory events in the underlying tissue.

#### Role of myelination

The presence of myelin along an axonal fiber is considered an indication of high excitability, because the nodes of Ranvier in between myelinated segments are known to contain a high density of sodium channels (Freeman et al. 2016). Hence, when estimating probability of activation due to stimulation, we included the effect of myelin by assuming a significantly higher activation threshold for unmyelinated fibers, compared to segments with nodes of Ranvier and axonal initial segments (see Methods: Computing the averaged probability of activation). Furthermore, non-uniform distribution of myelin can affect overall response to stimulation. Recently, an experimental study (Tomassy et al. 2014) of layer II/III pyramidal neurons revealed complex intermittent myelination patterns, where myelinated segments alternate with long unmyelinated paths. To reveal the possible impact of myelination on the activation probability of pyramidal neurons, we performed additional analysis. The leftmost column in Fig. 5 compares activation probabilities for myelinated and unmyelinated pyramidal neurons. layer II/III PYs showed no significant difference in activation probabilities due to myelination. In contrast, PYs from layer IV showed a drastic difference: with 80% activation in the presence of myelin and almost null activation probability for unmyelinated fibers.

Hence, in general, we found that cells with somas (and axonal initial segment) close enough to the superficial stimulation electrode and vertically oriented axonal arbors (like PYs in layer II/III) are likely to not see a great loss of activation probability if they lack myelination. In fact, our method shows that myelination plays a strong role in promoting cell excitability for cells which have axonal initial segments in deeper layers, further away from the current source (like PYs in layer IV), because they would rely more strongly on action potential in distal axonal fibers in the superficial layers to be generated by the input. The principle that unmyelinated fibers can be activated despite their lower excitability as long as they are placed closed enough to the current source applies also to interneurons. In fact, Bitufted cells and layer I interneurons (which are unmyelinated) show moderate activation probabilities in supragranular layers but no activation for deeper layers. Martinotti cells, although unmyelinated, do not show strong difference in their activation probability across cell layers. This is due to their specific shape, characterized by extensive axonal arborization in the upper layers.

#### Type and magnitude of stimulation control cell activation probability

In our estimates, the probability of cell activation depends directly on the fraction of axonal arborization which has activation function above threshold (length of the trigger area). In turn, this length depends on the amount of overall stimulation current delivered by the electrode (I). Intuitively, the larger the current magnitude, the longer the trigger area (Fig. 4), and hence the higher the spiking probability. However, it is less clear how changing the stimulation polarity would affect the likelihood of cell activation (Fig. 4), since different types of stimulation have completely different effect on axonal fibers: below the electrode, anodal current depolarizes vertical fibers and hyperpolarizes horizontal fibers, while cathodal current has the opposite effect (Fig. 1e,f). Therefore, we next asked how the probabilities of cell activation (Fig. 5) depend on the amplitude and polarity of the stimulating current I, and whether this dependence is similar across different cell types with different anatomies (Fig 2B). We applied our method to estimate activation probability (Fig 5) when the electrode delivered different amounts of total current I. We estimated this probability in a volume nearby the stimulating electrode, and repeated each estimate for the case of anodal and cathodal stimulation separately.

**Fig.6.**
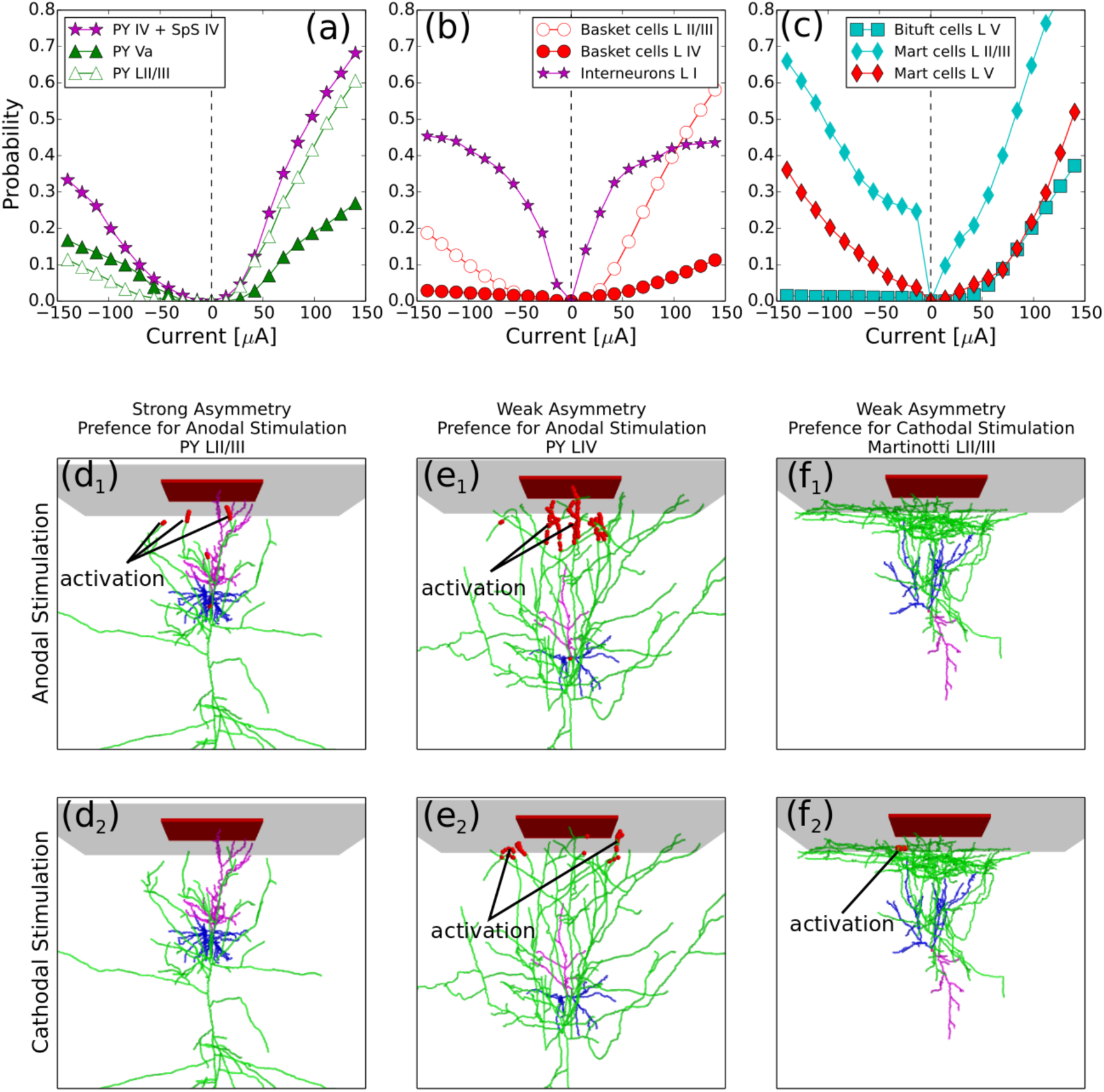
Different cell types have distinct preferences for stimulation type (anodal or cathodal) **(a-c)** Dependence of the activation probability on the net electrode current *I* for excitatory cells **(a)**, basket cells **(b)**, Martinotti and bi-tufted cells **(c)**. **(d_1,2_)** Anodal stimulation **(d_1_)** activates pyramidal cells LII/III more effectively than cathodal stimulation **(d_2_)**. **(e_1,2_)** Pyramidal LIV/V and spiny stellate cells have no preference for any type of stimulation. Because of the rich axonal arborization in supragranular layers, both types of stimulation provide large activation area. **(f_1,2_)** Cathodal current is more effective in activation of non-myelinated horizontal axons of Martinotti cells in supragranular layers.

Figure 6(a-c) shows how the overall activation probability changes as a function of the net current of stimulation (the total current used for Fig. 5 was 100 μA) for different cell types. All cells can be divided into three main categories based on their response types to the current: cells which respond more strongly to anodal stimulation, cells with mild preference for responding to anodal stimulation, and cells which respond more strongly to cathodal stimulation. In fact, a cell position across cortical layers and shape of its axonal arborization defines its preference to stimulation type.

#### Cells with strong preference for anodal stimulation

Pyramidal neurons, basket cells from layer II/III and bi-tufted cells constitute the first class, which is characterized by strong preference for anodal stimulation. The probability dependence on the current was very asymmetric for this class of cells (Fig.1(a)) due to much higher probabilities for positive *I > 0* current (anodal stimulation). Figure 6(d_1,2_) shows examples of the trigger areas (red markers) for a pyramidal cell from LII/III exposed to anodal vs cathodal stimulation. As can be seen in Fig 6d_1_, anodal stimulation activated several vertically oriented branches close to the soma. In fact, in general anodal stimulation depolarizes vertical fibers (Fig. 1e, f). In contrast, the same magnitude of cathodal stimulation was not able to produce any trigger area in this example (Fig.6d_2_), because cathodal stimulation is not effective at depolarizing vertical fibers.

#### Cells with weak preference for anodal stimulation

The second class includes cells which showed a less asymmetric relation between probability and current (Fig.6(a,b)) and did not show very strong preference for a type of stimulation. In this group we found spiny stellate cells, PYs from layer IV and slender-tufted PYs from layer. As a representative example of this case, we show in Fig. 6e_1,2_ the trigger area of a LIV PY cell exposed to anodal vs cathodal stimulation. This cell has an extensive axonal arborization in layers I and II, containing a large number of variously oriented fibers. This arborization with no dominant directions results in a response profile which cannot differentiate between anodal and cathodal current input, because the lengths of the trigger area created by the different types of stimulation are similar (even if specific fibers which cross the threshold are different). This is evident in Fig 6e_1,2_, where the red markers (trigger area) are different in the two panels, but cover similar amount of the axonal arbor. Similar arborization profiles, with no specific dominant orientation among the axonal fibers, are typical of all cell types included in this category (Fig 2), and hence result in a similar lack of selectivity of these cells for anodal or cathodal stimulation.

#### Cells with preference for cathodal stimulation

Our analysis indicates that Martinotti cells are the one cell type in this group which shows, at relatively small magnitudes (|*I*| ≾ 75 μ*A*), a strong preference for cathodal stimulation, (Fig. 6c). When, however, the stimulation current is strong, they respond to either stimulation type. The strong response at low magnitudes for cathodal current in MCs is driven by their peculiar arborization. These cells are characterized by a large number of horizontally oriented fibers in layer I (Fig.2B, Fig.6f_1,2_), which are likely to get activated in the presence of cathodal stimulation (Fig.1e). Hence, even small amounts of cathodal current, not capable of reaching deep layers, still induce enough trigger area in MCs axonal arbor (note that since MCs are unmyelinated, they do not require a large trigger area for activation). In contrast, for stronger current magnitudes, a similar probability of MCs activation is induced by anodal or cathodal stimulation. In fact, larger currents reach deeper layers, where the overall axonal arborization of MCs also includes vertical fibers. This results in a chance for anodal stimulation to trigger activation, and hence to have effects comparable to cathodal stimulation of the same magnitude.

Interneurons from LI do not clearly fit in one of the categories described above. In particular these cells have quite symmetric dependence of activation probability on curent. This is likely because LI interneurons locate close to the electrode and can be directly activated (through axonal hillock) by anodal stimulation. From the other hand, layer I interneurons are characterized by horizontally oriented axonal fibers, much similar to MCs. Hence, their activation probability depends on cathodal current in a very similar trend as MCs.

#### Network simulations predict optimal currents to trigger population response and strong difference for anodal and cathodal stimulations

In what is above we focused on direct activation of individual cortical neurons by the stimulation current, but ignored that these neurons can be synaptically connected. In order to predict an overall network response, we designed a computational model of the cortical column receiving surface cortical stimulation.

The cortical model included four types of cortical neurons organized within a multi-layer and experimentally verified connectivity structure. Namely, the model contained three distinct layers (layer II/III, layer IV and layer V) and each layer included excitatory neurons (PYs and SCs) and inhibitory interneurons (BCs and MCs). We omitted the other interneuron types because of their low density (Markram *et al.* 2004). The neurons in the model had probabilistic synaptic connections, organized according to canonical microcircuit architecture (Douglas *et al.* 1989; Thomson *et al.* 2002; da Costa and Martin 2010; Defelipe *et al.* 2012). The overall connectivity of the network is shown in Fig 7A: (a) all excitatory cells within each layer had recurrent excitatory connections, (b) PYs and SCc from layers IV and Va had strong projections to PYs from layers II/III, (c) BCs formed strong inhibitory connections to excitatory neurons within their own layers, and (d) MCs from layers IV and V made cross-layer inhibitory connections specifically targeting excitatory cells in layer IV. Two electrophysiological classes of neurons were used in the model: regular spiking neurons to represent PYs, SCs and MCs and fast spiking interneurons to represent BCs (Fig.7B). All inhibitory interneurons had lower leak current, which resulted in a higher responsiveness of these cells in comparison to excitatory neurons (Povysheva *et al.* 2006). Without input, the neurons were assumed to remain silent. For the first 1 ms, all cells were activated according to the probabilities estimated from our analysis (Figs 5, 6)), which was tailored to a stimulation duration of 200μs (Tehovnik *et al.* 2006). (Equations, parameters and details of the rationale followed in the design of our computational model are reported in Methods, Computational model of the canonical cortical circuit: rationale and parameters.)

We tested the model for different types (anodal vs cathodal) and magnitudes of current stimulation. When current magnitudes were low, the probability of direct activation was low (Fig.6(a-c)). In the network, it led to the sparse spiking in few pyramidal cells and inhibitory interneurons. However, no lasting population response was evoked (data not shown). Fig. 7c shows representative examples of the network response to moderate and strong magnitudes of surface current stimulation: the three panels compare the activity elicited in cells by different types of stimulation (cathodal in left panel, and progressively larger anodal stimulation in the two right panels).

The left panel in Fig.7c shows the network response to moderate cathodal stimulation. In our estimates, we found that cathodal stimulation triggered spiking in a smaller number of PY in LII/III, but activate a larger number of MCs and layer I interneurons in comparison to anodal stimulation. Hence, we hypothesize that cathodal stimulation can recruit strong inhibition in the tissue below the electrode, and produce weaker overall response. When tested in simulations, this hypothesis held true. In fact, cathodal stimulation evoked spikes in a large number of MCs in all layers, including LIV and V (red dots) where they constitute a large fraction of all interneurons (Markram *et al.* 2004). Note that MCs from infragranular layers specifically target excitatory cells in LIV (Wang *et al.* 2004). Hence, the effective recruitment of inhibition and relatively small activation probability of LII/III PYs resulted in a weak and sparse response of excitatory cells in our model column (left panel in Fig.7c).

The middle panel of Fig 7c shows that anodal stimulation of the intermediate intensity could trigger network activity across the layers for a considerable period after stimulus offset. In fact, stimulation triggered abundant spiking in mutually excitatory PYs and SCs in LIV, and some spikes in PYs in LII/III. In turn, the connectivity within LIV created reverberating excitatory activity that promoted a strong local network response, which projected onto LII/III PYs, eliciting a strong population response within L II/III. While activity of PYs in layers IV and II/III did outlast the duration of current input, it also recruited feedback inhibition from BCs and MCs, leading to termination of the response activity. Our simulations show that anodal currents of moderate magnitudes can induce a functional response beyond the stimulation duration and location. In fact, firing of LII/III PYs had the potential to reach other columns and deep layers PYs, which could then transmit to other areas the activity elicited in the network by the brief surface stimulus.

**Fig.7.**
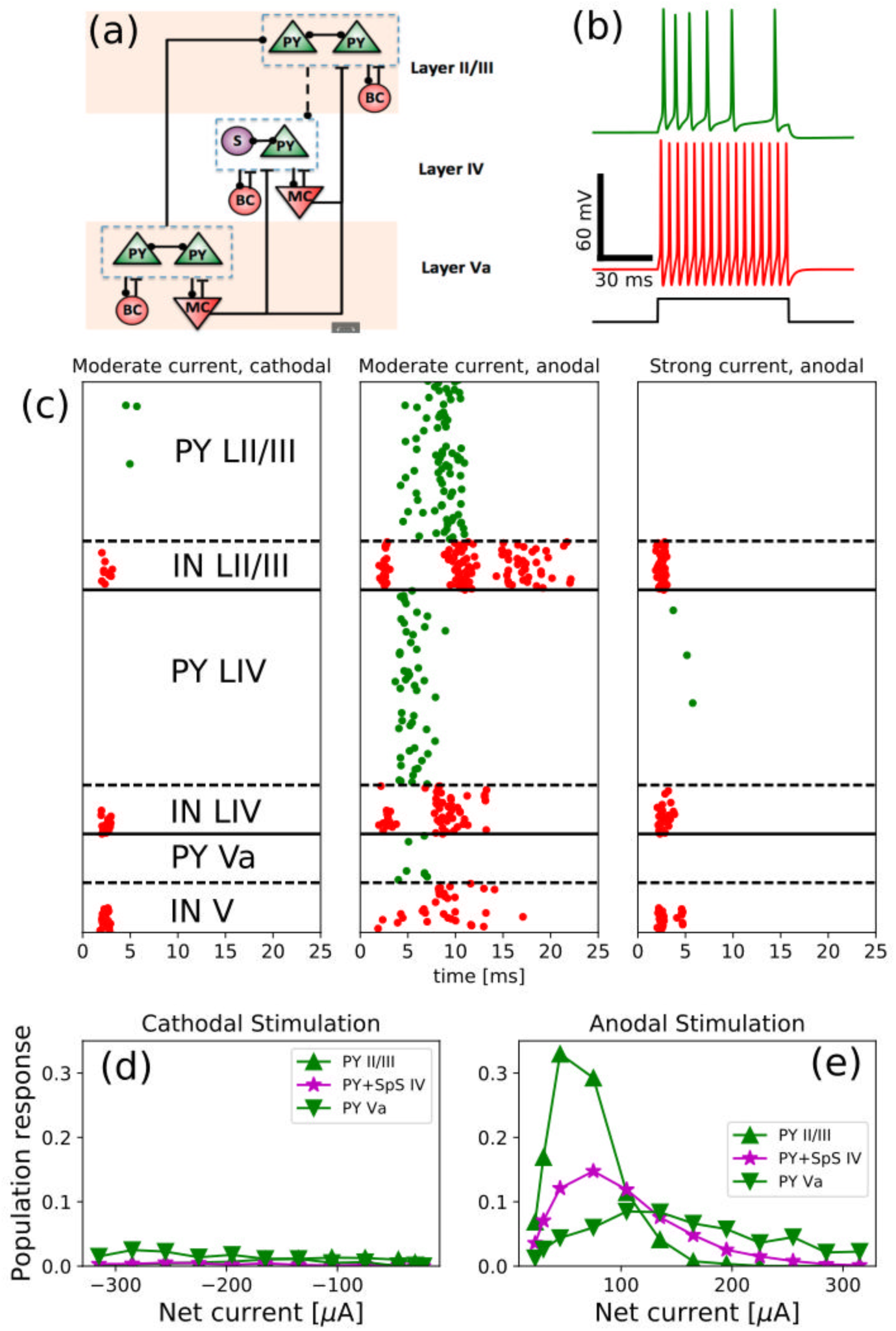
Numerical simulations predict that feedback inhibition controls response properties. **(a):** Schematic representation of the network model structure, which consists of 3 types of cells, located in 3 different layers (canonical circuit). PY stands for pyramidal neuron, SpS-spiny stellate cell, BC-basket cells, MC-Martinotti cells. Lines with circles denote excitatory AMPA connections (solid-strong, dashed-weak), whereas bars denote inhibitory GABA connections. **(b):** Two electrophysiological classes of neurons were used in our simulations: top voltage trace (green) corresponds to regular spiking neurons (used for pyramidal, spiny stellate cells and Martinotti cells) and bottom voltage trace demonstrates activity of fast spiking interneurons (used for basket cells). **(c):** Spike raster plots exhibit network activity for cathodal (left panel) and anodal (two right panels) stimulations. The cells were activated during first 1 ms of simulation according to activation probability (see text for details). Green dots-PY and SpS cell spikes, red dots-interneuron spikes. Left panel shows weak response to cathodal stimulation (-100 μA). Middle panel shows response to moderate anodal stimulation (75 μA), which induced a large population response. The right panel shows response to a strong positive current (300 μA), which activated a large number of basket cells in layer II/III and Martinotti cells in all layers, which prevented the activation of excitatory cells in layers II-IV. **(d,e):** Population responses as a function of net electrode current for layer II-IV excitatory cells.

In the case of strong anodal stimulation, our estimates showed a high probability of activation for most cell types (Fig 6a-c). Once connected, our simulations predicted a network response dominated by inhibitory activity (Fig. 7C, right panel). In fact, current stimulation activated both interneurons and excitatory cells, but the larger input resistance and higher responsiveness of interneurons compared to PYs and SCs resulted in the activation of these inhibitory neurons before the spikes in excitatory cells. Because of the large number of interneuron spikes evoked, synaptic inhibition on PYs and SCs could overcome the excitatory drive due to stimulation, and effectively stopped them from firing. Because of the lack of directly recruited inhibition in layer Va, slender-tufted PYs were activated by stimulation, but their relatively low density meant that they could only deliver a small amount of excitation to the highly inhibited upper layers.

While Fig 7c shows few characteristic examples, panels d and e in Fig. 7 summarize the findings across simulations with many current magnitudes. These plots show the average population response (computed as a number of spikes divided by the total population number) for excitatory cells as a function of applied stimulation current *I* (cathodal in panel d and anodal in panel e). For low current, probability of direct activation was low in both anodal and cathodal stimulations, whereas for large current, inhibitory activity suppressed spiking in PYs and SCs. Importantly, for excitatory cells and anodal stimulation there was an optimal range of stimulation current magnitudes where population response was highest (consistent with our description of network dynamics in Fig 7c, middle panel). In fact, the input evoked moderate amount of excitatory spiking amplified by recurrent excitation, which led to the high overall response in the cortical column.

## Discussion

In this study we estimate the probability of activation for different cell types and in different cortical layers when exposed to external electrical stimulation. We present the case of a finite size electrode placed on the cortical surface. The approach consists of 4 main steps: first, estimate the electric field potential in the tissue; second, define the ‘axonal-electrical receptive field’ based on reconstructions of different cell types across layers; third, the estimate of cells’ spiking probability based on the activation function in axonal elements; finally, predict the network response to stimulation in a minimal model of a cortical column based on the spiking probability estimates. Our study predicts that short-lived superficial stimulation with a single electrode source has ability to trigger spiking in layer IV pyramidal cells, and to evoke network activity that could persist for hundreds of milliseconds. It further predicts a much higher spiking response to anodal stimulation compared to cathodal one, as well as existence of the optimal stimulation intensity, capable to induce a maximal response in a population of cortical cells.

### Relevance of our findings to existing stimulation protocols

Recent advances in techniques of multisite cortical stimulation (Lewis et al. 2015), aimed to restoring damaged brain operations like movement, sensation, perception, memory storage and retrieval, underscore the need for better understanding the effects of such stimulation on individual neurons and synaptic connections. Our study predicts that local electrical stimulation may elicit activity under a superficial electrode sufficient to enable signal transmission without trumping the physiology of the network. Indeed, in our study, stimulation within an intermediate range of positive input currents triggered network response that survived the end of the stimulus, and thus can potentially be similar to the physiological processing in the stimulated tissue. The last is important if the stimulation targets to induce physiologically relevant persistent changes in brain tissue. For example, inducing synaptic plasticity is necessary for stimulation to be successful as an intervention to restore memories (Stagg and Nitsche 2011; Kuo and Nitsche 2012), promote the recovery of a cortical area lost functions (Jackson et al. 2016; Malerba et al. 2017), or repress hyper-excitability of a portion of tissue (Liebetanz et al. 2006). But blanket plasticity evoked by the cell spiking that is directly triggered by external current is not likely to match existing synaptic patterns and will not have any meaningful constructive impact on a system of careful counter-balances like the brain. In contrast, stimulating in the range of currents that elicit network-driven activity that continues when the stimulus is removed provides a better chance for stimulation to be changing only a selected and physiologically meaningful subset of synapses. In other words, there is a range of current values in which stimulation can be used as a sophisticated and detailed intervention rather than a blunt hammer, and such range can be found combining theoretical estimates of the current density and reconstructed anatomy.

Our work focuses on the effects of a single brief isolated pulse, in contrast to repetitive trains or longer-lasting modulation, which are typically used in the clinical setting (such as in cortical stimulation mapping, neurorehabilitation with tDCS, etc). Also clinical stimulation is often bipolar (between two electrodes) and biphasic, with anodal and cathodal current balanced in order to prevent irreversible Faradic currents (Merrill et al. 2005; Cogan et al. 2016). Furthermore, our charge density is ~0.06μC/cm2, far below levels commonly used in clinical stimulation. Of course, activation thresholds are probably higher in humans due to thicker pia and cortex, but detailed neuronal reconstructions are not available to model the effects of these anatomical species differences. Thus, our results, as presented, cannot be applied directly to the usual clinical context, and future studies are necessary to connect our findings to the clinical realm. Specifically, in the case of weak and long time scale current waveforms (like tDCS) the stimulation is capable of modulating cell properties, and our method would need to incorporate estimates of the dendritic dynamics in the presence of extracellular electric fields (for example using the equivalent cylinder models (Rall 1962; Tranchina and Nicholson 1986). Also, trains of stimulation pulses (such as in cortical mapping) will have effects beyond the simple summation of the effects of single pulses, due to intrinsic cell dynamics and synaptic interactions, and both dendritic and axonal dynamics will need to be included in the spiking probability estimates (using for example multi-compartmental models (Berzhanskaya et al. 2013). However, as demonstrated experimentally (Tehovnik *et al.* 2006), the threshold for evoking action potentials by stimulation of the type we consider is by far the lowest at the nodes of Ranvier, and next lowest at the axon hillock. While transmembrane currents can be conducted intracellularly from the dendritic tree to the axon hillock, they are greatly delayed and attenuated, and their effects are minor compared to the direct stimulation of these elements.

Our results suggest that superficial anodal stimulation is more effective than cathodal at cell activation. Clinically, with respect to cathodal vs anodal stimulation effects, the evidence is somewhat mixed. Previous work by (Hern et al. 1962), among others found that surface-anodal stimulation had lower threshold for activation of corticofugal fibers in the Baboon’s motor cortex. In Pollen’s classic studies (Pollen 1977) recording units in cat visual cortex in response to stimulation of the overlying surface, he noted that different cells had lower threshold to either cathodal or anodal currents. A possible explanation can be found in the modeling study (Manola et al. 2007) which found that the although surface-anodal current would preferentially activate vertically oriented elements directly beneath the electrode (as in our study), elements in the sulci would activated by cathodal current, and these effects interacted with the precise location of the electrodes with respect to gyral crowns and whether bipolar stimulation was used.

### Limitations of the approach

Experimental studies over past decades found that electrical microstimulation directly activates axon initial segments and nodes of Ranvier (Porter 1963; Gustafsson and Jankowska 1976; Swadlow 1992; Nowak and Bullier 1998, 1998; Rattay 1999), which are the most excitable elements due to the high concentration of sodium channels (Catterall 1981). Thus, the estimation of direct activation of cortical cells in our study was based on calculation of the activation function along the axonal segments. This approach takes into account orientation and thickness of axons, and also considers their myelination properties. We defined a threshold for the effective stimulation current in an axonal segment (given by the activation function) by direct comparison to the experimental data: in turn, this threshold enabled the calculation of cell activation probability. In our analysis, we applied the same method to excitatory and inhibitory populations, which implies that all parameters of our estimation scheme (such as threshold for activation function) are the same for all cells. Since BCs have higher input resistance compared to PYs, it is possible that their experimental threshold could be lower than our estimate, which should not, however, affect our conclusions: in fact, the suppression of excitatory firing at high input current values would still hold, since it hinges on BCs spiking before PYs (which would only be enhanced by lowering the threshold).

Another potential limitation of our approach is that in our estimates we considered a homogeneous tissue. Inhomogeneity in the tissue would affect the trigger area along axons, and could change activation properties in each particular cell. Since the main source of such inhomogeneity is other neurons or glia, its effect is expected to be stronger on the deep layer neurons. While the exact effects of inhomogeneity still need to be explored, we believe that the specifics of our approach can mitigate such effects. Indeed, our estimate averages across cell rotations, shifts and multiple different reconstructions to compute a probability of spiking. This implies that the effect of local changes in the tissue (driven by inhomogeneity) would likely affect the final average probability only marginally. Hence, we do not expect dramatically different estimates for cell activations to emerge from finer estimates of tissue properties.

The connectivity within our network model was based on the canonical model, which captures the main picture of the information flow across cortical layers (Douglas *et al.* 1989; Thomson *et al.* 2002; da Costa and Martin 2010; Defelipe *et al.* 2012), yet overlooks finer properties like the descending projections from excitatory to inhibitory neurons (Thomson *et al.* 2002), the activity of less frequently observed interneurons (bi-tufted, neuroglia form, etc.), the fine variability of pyramidal cells within layer II/III PYs and their projections (Shepherd and Svoboda 2005; Staiger *et al.* 2015). Also, the network model predictions can be extended toward multi-compartment neuronal models (Bonjean et al. 2012), which can take into account finer structure of inhibitory targeting (e.g. tuft vs soma-targeting interneurons) (Markram *et al.* 2004) and explicitly model propagation of orthodromic and antidromic spikes. The exact and detailed aspects of inter-layer connections, which update the canonical model (Dhruv 2015) are beyond the scope of our work, which focuses on calculating the probability of direct activation of cortical cells as a result of external electrical stimulation.

Our estimates are based on activation function and do not consider the voltage dynamics within axonal trees. However, theoretical work addressing axonal dynamics through multi-compartmental modeling (Rattay 1999) has shown that for moderate currents the activation function correlates with voltage dynamics, and predicts the activation sites within axonal elements. For very strong currents, areas along axonal arborization could be inactivated, blocking propagation of action potential along the axon (Rattay and Aberham 1993). In this case, our estimate could not apply. However, the blocking phenomenon arises only for relatively strong stimulation currents and typically affects elements, which are very close to the electrode (Rattay 1987; Rattay and Aberham 1993). Hence, we expect that for small stimulation currents (considered in our study) and superficial electrode configuration, the blocking phenomenon does not qualitatively change the results of our study. Overall, the details of axonal propagation dynamics are not essential to our method, since it focuses on average estimates of response for a cell type, rather than an account of stimulation-induced dynamics in a specific cell.

### Generalization of the method

While we applied our analysis to the microstimulation protocol by a single superficial electrode (Ha *et al.* 2016), our strategy can be directly generalized to a number of more complex settings by using linear summation and adjusting the calculations for the current density: multiple electrodes (as in ENIAC (Ha *et al.* 2016)), in-depth stimulation (as in epilepsy (Nagaraj *et al.* 2015)), non-circular electrode plates (as in DCS (Hummel and Cohen 2006)) and bi-polar electrodes (as in DBS (Blumenfeld and Brontë-Stewart 2015; Baizabal-Carvallo and Alonso-Juarez 2016; Papageorgiou *et al.* 2016)) can all be accounted for. Also, we assumed a generic cortical tissue volume but the same idea can be applied to the tissue different from a canonical cortical column (e.g., hippocampus, geniculate nuclei, damaged tissue after traumatic brain injury or stroke) as long as enough data on reconstructed cells are available.

Since different stimulation protocols use different current waveforms, it is important to note that this approach can be generalized to the other stimulation protocols as long as an activation threshold has been experimentally measured. The type of activity elicited by stimulation can also be expanded. Our method is presented in the context of evaluating activation mediated by axonal spikes, an effect relevant for fast and strong stimulation protocols (ENIAC, DBS), but other types of stimulation could be focused on triggering subthreshold effects, such as voltage polarization at somas compared to dendrites (tDCS) (Bikson et al. 2012). To adapt the method to account for these sub-threshold effects, one would have to embed the reconstructed cells in the electric field and consider a probability of depolarization/hyperpolarization (Ascoli *et al.* 2007), taking into account the specific orientation of each element of the reconstructed cell compared to the direction of the current field.

### Conclusions

We introduced a generic approach to estimate probabilities of cell activation in response to external stimulation, and applied it to make testable predictions regarding effects of superficial electrical microstimulation of a canonical cortical circuit. Ongoing rapid extension of the available data on the reconstructed neuronal anatomy will make possible to increase the precision of our analysis and to apply it to other brain regions and species. Our study shows an example of how such basic knowledge can play directly into designing models that further our understanding of brain function and its interaction with devices.

## Acknowledgements

The authors would like to thank Dr. Giri Krishnan, Matthew Choinski, Dr. Nikolai Rulkov, Dr. Bruce Mc Naughton, Dr. Gert Cauwenberghs, Dr. Terry Sejnowski and Dr. Patrick Mercier for insightful discussions. This work was supported by UC MRPI (MR-15-328909) and ONR (MURI: N000141310672).

